# European Vintage tomatoes galore: a result of farmers combinatorial assorting/swapping of a few diversity rich loci

**DOI:** 10.1101/2021.10.26.465840

**Authors:** Jose Blanca, Clara Pons, Javier Montero-Pau, David Sanchez-Matarredona, Peio Ziarsolo, Lilian Fontanet, Josef Fisher, Mariola Plazas, Joan Casals, Jose Luis Rambla, Alessandro Riccini, Samuela Pombarelli, Alessandra Ruggiero, Maria Sulli, Stephania Grillo, Angelos Kanellis, Giovanni Giuliano, Richard Finkers, Maria Cammareri, Silvana Grandillo, Andrea Mazzucato, Mathilde Causse, Maria José Díez, Jaime Prohens, Dani Zamir, Joaquin Cañizares, Antonio Jose Monforte, Antonio Granell

**Affiliations:** Instituto de Conservación y Mejora de la Agrodiversidad Valenciana (COMAV-UPV), Universitat Politècnica de València, València, Spain; Instituto de Biología Molecular y Celular de Plantas (IBMCP). Consejo Superior de Investigaciones Científicas (CSIC), Universitat Politècnica de València, València, Spain; INRAE, UR1052, Génétique et Amélioration des Fruits et Légumes, 67 Allée des Chênes, Centre de Recherche PACA, Domaine Saint Maurice, CS60094, Montfavet, 84143, France; Hebrew Univ Jerusalem, Robert H Smith Inst Plant Sci & Genet Agr, Rehovot, Israel; Department of Agri-Food Engineering and Biotechnology/Miquel Agustí Foundation, UPC- BarcelonaTech, Campus Baix Llobregat, Esteve Terrades, 8, 08860 Castelldefels, Spain; Department of Agriculture and Forest Sciences (DAFNE), Università degli Studi della Tuscia, Viterbo, Italy; Institute of Biosciences and BioResources (IBBR), National Research Council of Italy (CNR), Via Università 133, 80055 Portici, Italy; Italian National Agency for New Technologies, Energy and Sustainable Economic Development (ENEA), Casaccia Research Centre, Rome, Italy; Group of Biotechnology of Pharmaceutical Plants, Laboratory of Pharmacognosy, Department of Pharmaceutical Sciences, Aristotle University of Thessaloniki, 54124 Thessaloniki, Greece; Wageningen Univ & Res, Plant Breeding, POB 386, NL-6700 AJ Wageningen, Netherlands

**Author notes:** Corresponding authors Antonio Jose Monforte, Antonio Granell. M.J. Díez. J. Cañizares. J. M.-G. Current address: Cavanilles Institute of Biodiversity and Evolutionary Biology (ICBiBE), Universitat de València, 46022, Valencia, Spain. L. M. Current address: HM Clause, Portes-lès-Valence, France.

**Keywords:** Crop evolution, diversification, selection, genotyping by sequencing, GWAS, SNP, fruit morphology

## Abstract

A comprehensive collection of 1,254 tomato accessions corresponding to European heirlooms and landraces, together with modern varieties, early domesticates and wild relatives, were analyzed by genotyping by sequencing. A continuous genetic gradient between the vintage and modern varieties was observed. European vintage tomatoes displayed very low genetic diversity, with only 298 loci out of 64,943 variants being polymorphic at the 95% threshold. European vintage tomatoes could be classified in several genetic groups. Two main clusters consisting of Spanish and Italian accessions showed a higher genetic diversity than the rest varieties, suggesting that these regions might be independent secondary centers of diversity and with a different history. Other varieties seem to be the result of a more recent complex pattern of migrations and hybridizations among the European regions. Several polymorphic loci were associated in a GWAS with fruit morphological traits in the European vintage collection, and the corresponding alleles were found to contribute to the distinctive phenotypic characteristic of the genetic varietal groups. The few highly polymorphic loci associated with morphological traits in an otherwise diversity-poor genome suggests a history of balancing selection, in which tomato farmers maintained the morphological variation by applying a high selective pressure within different varietal types.

**Highlight:** The high phenotypic diversity observed among European vintage varieties was created by traditional farmers by combining very few polymorphic loci subjected to balancing selection.

## Introduction

The widespread tomato crop (*Solanum lycopersicum* L. var. *lycopersicum*; SLL) originated in Mesoamerica in a region corresponding to today’s Mexico as a result of the *S. lycopersicum* L. *var. cerasiforme* (SLC) (Blanca *et al*., 2012; Blanca *et al*., 2015; Razifard *et al*., 2020). Tomato was later brought to Europe, and the Italian botanist Mattioli in 1544 already described varieties with flat, round and segmented fruit types (McCue 1952). This indicated that tomato had probably arrived to Europe in different shapes from America (Luckwill, 1943; Sanfuentes-Echevarria 2006; Sahagún 1577). Tomato was not immediately adopted for consumption by Europeans, as it was considered at different times and regions as: poisonous, aphrodisiac, ornamental, valuable for sauces and soups, miracle cure and, finally, a fresh salad ingredient (Harvey 2004). It was only as late as the mid-19th century that the tomato became a regular component of the diet in Britain and North America (Harvey 2004). On the contrary, the tomato was better received, extensively cultivated, and consumed as food by the 18th century in Southern Europe, which therefore could have become a secondary center of diversity (Boswell 1937; Bauchet and Causse 2012). As a result of this long tradition of use a large number of traditional varieties are currently available along the Mediterranean basin showing an impressive phenotypic diversity in terms of fruit appearance, adaptation to local conditions and culinary use. Despite the interest for unveiling the population history and the processes that gave rise to the domestication of tomato (Blanca *et al*., 2015; Razifard *et al*., 2020), there are yet no detailed genetic analyses of the diversification history of the European traditional tomato varieties.

The extent and type of the molecular variation in the tomato clade has been extensively analyzed in previous studies. The first molecular studies, carried out with isoenzymes, determined that the worldwide cultivated SLL was less variable than the wild *S. pimpinellifolium* (SP) and that the wild, feral and semi-domesticated *S. lycopersicum* var. *cerasiforme* (SLC) was genetically closer to SLL than to SP (Rick *et al*., 1974; Rick and Fobes 1975). A clear trend of diversity reduction was already observed at the species/subspecies level, probably due to bottlenecks associated with migrations and to the selection pressure imposed by humans during the early domestication stages and development of cultivars from SP to SLC, and lastly, to SLL, (Blanca *et al*., 2012, 2015, Razifard *et al*., 2020).

Despite this limited SLL diversity, several molecular studies have unveiled the worldwide genetic structure within SLL, dividing it into four major groups: processing and fresh market, cherry and vintage tomatoes (Williams and St. Clair 1993; Robbins *et al*., 2011; Sim *et al*., 2011; Casals *et al*., 2019). The first three groups correspond to modern tomato varieties created by breeders in the 20th century, characterized by their different culinary use and the introgression of wild species genes, mainly to increase disease resistance and also to develop new type of cultivars. Vintage cultivars are defined as those developed by traditional farmers by intuitive breeding and were cultivated (and some of them are still nowadays locally) before the advent of professional breeding. In this study, landraces, traditional and heirlooms are considered as synonymous of vintage. Park *et al*., (2004) found genetic differentiation between vintage and modern cultivars. A more comprehensive analysis using 7,720 SolCAP single nucleotide polymorphisms (SNP) from over 426 accessions confirmed the previously described fresh, processing, and vintage groups, at the same time finding two extra clusters located between SLL and SP that corresponded to cultivated and wild cherry tomatoes (Sim *et al*., 2012). Blanca *et al*., (2012; 2015) also obtained the fresh, processing, and vintage split and clarified the status of the cherry tomatoes: some of them were SLC from South America, Mesoamerica, and the subtropical regions, while others were modern cherry tomatoes obtained by hybridizing cultivated SLL with wild SP. *Blanca et al*., (2015), compared with a rarefaction analysis the genetic diversities of the different groups and found that vintage SLL and SLC from outside Peru and Ecuador had the lowest diversity, whereas Peruvian and Ecuadorian SP and SLC had much higher diversities.

The studies mentioned above differentiated the modern varieties from the vintage ones, but none of them found any structure within the vintage tomato group. García-Martínez *et al*., (2006) studied a collection of vintage Spanish cultivars belonging to the varietal groups “Muchamiel”, “Pera”, and “Moruno” with 19 microsatellite and amplified fragment length polymorphism markers and managed to differentiate the “Pera” type from the other two groups. *Mazzucato et al*., (2008) dissected a collection of 36 Italian vintage accessions by using 29 microsatellites, and Sacco *et al*., (2015) found differences between 61 Italian vintage varieties and 26 American ones. Current genomic sequencing technologies allow finding variable molecular markers even in very narrow genetic contexts. Thus, recently, Esposito *et al*., (2020), using double digest restriction-site associated DNA sequencing (ddRAD-seq), was able to obtain a sufficient number of SNPs to study the differentiation of a special type of vintage tomatoes cultivated in Spain and Italy, called “de penjar” or “da serbo”, characterized by their long shelf-life (LSL). Overall, “de penjar/da serbo” varieties tended to cluster together, showing certain genetic differentiations when compared with other vintage and modern cultivars, but some level of admixture was also found. These former studies were focused on a limited number of accessions from a narrow local diversity and therefore a broader view is clearly needed to better understand the history and relationships of the European vintage varieties.

In the present study, the genomes of 1,254 European tomato accessions collected from Southern European seed banks were partially sequenced by Genotyping by Sequencing (GBS, Elshire *et* al., 2011; Baird *et al*., 2008) to obtain genotypes for unbiased markers. Based on these, the genetic structure, diversity, and the association between the polymorphic loci with the morphological variation in that collection were analyzed to shed light on the history of the making up of the diverse vintage European tomatoes.

## Material and methods

### Materials

A total of 1,254 tomato accessions were analyzed in this study. One thousand forty four of these accessions are part of the collection of the EU TRADITOM project (www.traditom.eu). Seeds composing the TRADITOM collection were obtained from the genebanks of the Institute for the Conservation and Improvement of Valencian Agrodiversity at the Polytechnic University of Valencia (COMAV-UPV, Valencia, Spain), of the Balearic Island University (UIB, Mallorca, Spain), the Station d’Amelioration des Plantes Maraicheres of the French National Institute for Agricultural Research, (INRA, Montfavet, France), of the Department of Agriculture and Forest Sciences of the University of Tuscia (UNITUS, Viterbo, Italy), of Institute of Biosciences and Bioresources of the Italian National Council of Research (CNR-IBBR, Portici, Italy), of of the Agricultural Research Center of Macedonia and Thrace of the National Agricultural Research Foundation (GGB-NAGREF, Thessaloniki, Greece) and the seed collections of the Miquel Agustí Foundation of the Polytechnic University of Catalunya (FMA-UPC, Casteldefels, Spain), of BioEconomy of the Italian National Council of Research (CNR-IBE, Catania, Italy), of ARCA 2010 S.C.ar.l. (ARCA, Acerra, Italy), of the University of Reggio Calabria (UNIRC, Reggio Calabria, Italy), of the Robert H. Smith Faculty of Agriculture, Food and Environment of the Hebrew University of Jerusalem (HUJI-ARO, Rehovot, Israel). An additional set of 110 accessions were obtained from the COMAV genebank (http://www.upv.es/contenidos/BGCOMAV/indexi.html) that contained 10 wild accessions from the Galapagos Islands, one accession of each wild species *S. habrochaites, S. chmielewskii* and *S. peruvianum*, 36 *S. pimpinellifolium* accessions from Peru (SP) and North Ecuador (SP_NECu) and 52 *S. lycopersicum* var. *cerasiforme* (SLC) accessions, three modern and 20 SPxSL (*S. pimpinellifolium* x *S. lycopersicum* hybrids, corresponding to cherry cultivars and other crosses between the two species). Passport data can be found in Supplementary Table S1. The germplasm collection was extensively phenotyped in the TRADITOM project (Pons *et al*., 2017, and in preparation). The dataset corresponding to fruit morphology and color traits obtained at the HUJI-ARO trial was used and analyzed for this article (Supplementary Table S2).

### DNA extraction, library preparation and sequencing

Genomic DNA was isolated from young leaves of 5-10 seedlings per accession, using the DNeasy 96 Plant Mini Kit (Qiagen, Germany). Genotype-By-Sequencing (GBS) was performed following the procedure reported by Elshire (2011). Briefly, DNA was digested with the restriction enzyme *Ape*K I, barcoded libraries were prepared to track each accession and the DNA sequence corresponding to the region flanking the *Ape*K I site was obtained on an Illumina HiSeq 2000 platform by LGC Genomics GmbH (Berlin, Germany). Following the Variant Call Format standard, we used the term sample to refer to one genotyping experiment from one accession.

### Read mapping, SNP calling and SNP filtering

FastQC was used to evaluate the quality of the sequenced reads, and these were mapped against the *S. lycopersicum* genome build 2.5 (Sato *et al*., 2012) using BWA mem (Li 2013). After mapping, the PHRED quality of 3 aligned nucleotides from each read end was set to 0 in order to avoid possible false positives caused by misalignments (Li 2011). Mapping statistics were calculated with the samtools stats command (Li *et al*., 2009).

SNP calling was carried out by freebayes (Garrison and Marth 2012) with the following parameters: a minimum mapping quality of 57, 5 best alleles, 20 minimum base quality, 0.05 maximum mismatch read alignment rate, 10 minimum coverage, 2 minimum alternate allele count, and 0.2 minimum alternate fraction. To avoid regions in the reference genome with potential assembly problems, the Heinz 1706 reads used to build the reference genome were mapped against the reference assembly version SL2.50, a 50X mean coverage was obtained, and when any region had a coverage higher than 200X was removed from the SNP calling.

SNP and genotype processing were carried out by using the variation Python library located at https://github.com/JoseBlanca/variation. To create the tier1 SNP set to be used in the rest of the analyses, the genotypes with a quality lower than 5 were set to missing, and the variants with a SNP quality lower than 50, an observed heterozygosity higher than 0.1, and a call rate lower than 0.6 were filtered out. Moreover, in order to avoid false positives, only variants in which the minor allele was found in more than 2 samples were kept. This filtering was carried out with the “create_tier1.py” script. For some analyses, the pericentromeric regions, that seldom recombine, were removed as part of the heterochromatin. To locate the pericentromeric regions a piecewise regression analysis was applied to the relationship between the genetic distance and the physical positions of the markers of the EXPIM map (Sim *et al*., 2012). Regression analyses were done using the segmented R library (Muggeo 2003). The calculated pericentromeric regions were: chromosome 1, from 5488553 to 74024603, chromosome 2, from 0 to 30493730, chromosome 3, from 16493431 to 50407653, chromosome 4, from 7406888 to 50551374, chromosome 5, from 9881466 to 58473554, chromosome 6, from 3861081 to 33077717, chromosome 7, from 4056987 to 58629226, chromosome 8, from 4670213 to 54625578, chromosome 9, from 6225214 to 63773642, chromosome 10, from 3775719 to 55840828, chromosome 11, from 10947270 to 48379978, and chromosome 12, from 5879033 to 61255621.

### PCoA and genetic structure, Diversities and Linkage disequilibrium

The genetic structure and the division in subpopulations were determined by conducting a series of hierarchical Principal Coordinate Analyses (PCoA). The PCoAs were carried out with a subset of the variants after filtering. The variant filtering process was comprised of several steps. First, only the euchromatic variants were considered, and from those only the ones with a call rate lower than 0.95, also the ones in which the minor allele was present in less than 3 samples were removed. From the remaining variants, 2000 evenly distributed across the genome were selected. Furthermore, in order to avoid overrepresentation of large regions with complete linkage disequilibrium, when several consecutive variants had a correlation higher than 0.95, only one of them was kept. Finally, pairwise distances between samples were calculated (Kosman and Leonard 2005), and from the distance matrix, a PCoA (Krzanowski and Krzanowski 2000) was generated following the pycogent implementation. These methods were implemented in the do_pca.py script. Additionally, the genetic structure was also estimated with fastSTRUCTURE (Raj *et al*., 2014).

The observed and expected heterozygosity and the number of variants per genetic group were calculated considering only the variants variable in the samples involved in the analysis. The script that implemented these analyses is calc_diversities2.py. The allele spectrum figure was plotted by the script calc_maf_trends.py and the rarefaction curves by rarefaction_analysis.py.

The linkage disequilibrium (LD) was calculated between euchromatic markers with a major allele frequency lower than 0.98 following the Rogers and Huff method for loci with unknown phase (Rogers and Huff 2009).

### GWAS and allele frequencies

A heatmap plot that represents the major allele frequency in each group was generated according to a dendrogram by the method implemented in the Python seaborn library (https://seaborn.pydata.org/) and was plotted by the get_most_diverse_snps.py script.

A Genome-Wide Association Analysis (GWAS) was carried out with the Genesys R package (Gogarten *et al*., 2019) on the set of polymorphic variants (95% threshold). The quantitative characters were normalized by using the Box and Cox transformation implemented by the Python scipy library (https://www.scipy.org/). The character normality was checked with a qqplot plotted by the Python statsmodels library (Seabold and Perktold 2010). The correction for genetic structure was calculated with a Principal Component Analysis on the filtered variants implemented by the SNPRelate R library with a 0.3 linkage disequilibrium threshold (Zheng *et al*., 2012). The quantitative trait associations were tested with the Wald method, and the binomial ones by the Score one. To account for the multiple tests, a Bonferroni threshold was applied. The step-by-step implementation of the GWAS analysis can be analyzed in the gwas.py script.

### Genetic group distances

Two genetic distances among groups were calculated and compared: Nei and Dest (Peakall and Smouse 2006; 2012). They were implemented by the Python variation library and the cacl_pop_dists.py script. From those distances both a neighbor joining tree and a split network were calculated using SplitsTree (Huson and Bryant 2006).

## Results

### High through-put genotyping of a European vintage tomato collection

To genetically characterize vintage European tomatoes, a total of 1,254 tomato accessions were used (Supplementary Table S1). That set included an extensive representation of the extant European vintage tomato variability constituted by 506 accessions from Spain, 305 from Italy, 203 from Greece, 96 from France, and 58 from other origins, with 25 modern commercial cultivars, 39 SP and 22 SLC accessions (the two last ones of American origin) used as references. A total of 3,700 million reads with a mean phred quality of 33.5 were obtained after genotyping-by-sequencing, providing an average of 2.9 million reads per sample. Out of those, 99.0% were successfully mapped to the tomato reference genome (v2.50), but only 55.9% were kept after applying the MAPQ filter with a 57 threshold. These reads were mostly properly paired (96.1%). Of all of the genomic positions that comprise the reference genome, 0.79% had a per sample average sequencing coverage higher than 5X, 0.46% higher than 10X and 0.21% higher than 20X. The complete sequencing and mapping statistics for all samples are available in Supplementary Table S3 and the number of positions per megabase with more than 5 reads in at least 90% of the samples is represented in Supplementary fig 1. Finally, 448,121 variants were called by freebayes, and after filtering them, a working dataset of 64,943 variants was created.

### Genetically defining true European vintage tomatoes and its relationship with American relatives

To genetically position the European tomato collection relative to South and Mesoamerican germplasms, which represent early domestication and improvement steps (Blanca *et al*., 2015), the observed variability of European tomato was analyzed together with SP, SLC, SLxSP hybrids and a sample of modern cultivars including modern fresh and processing cultivars. A series of PCoAs (Fig. 1 and 2) was performed comparing vintage and modern vintage collections. The genetic classification based on these PCoAs can be found in Supplementary Table S1 under the header rank1 classification.

**Fig. 1.**
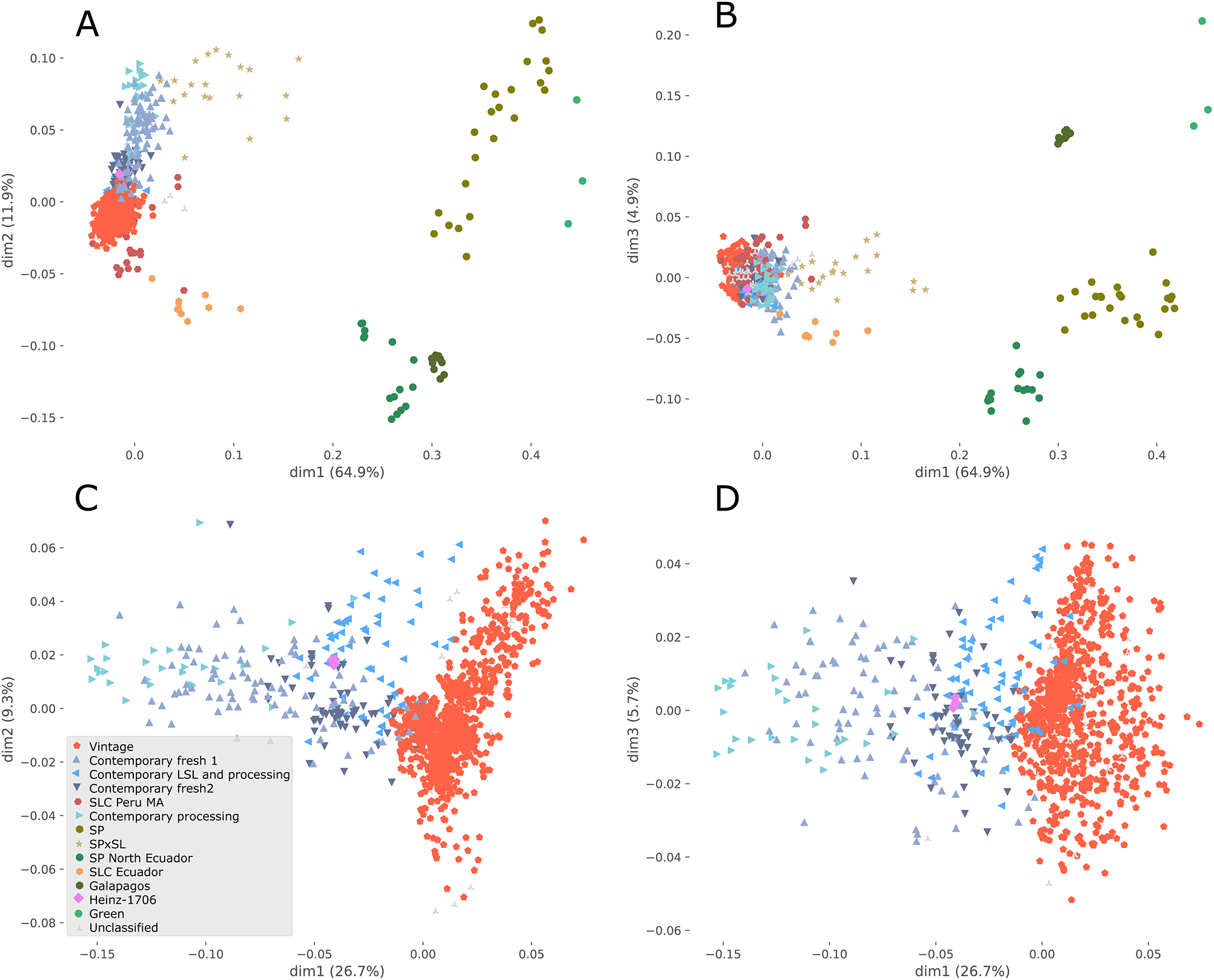
Principal Coordinate Analysis (PCoA) including cultivated tomato (*Solanum lycopersicum* var. *lycopersicum*, SLL): vintage European tomato, modern cultivars with different culinary use (fresh, processing and long shelf life, lsl), *S. lycopersicum* var. cerasiforme (SLC) from different origin [Peru, Mesoamerica (MA) Ecuador (Ecu)], together with several American wild relatives: *S. pimpinellifolium* (SP), *S. cheesmaniae, S. galapagense* (Galápagos), *S. peruvianum, S. chmielewskii* and *S. habrochaites* (green) and SPxSL hybrids. The modern cultivar Heinz1706 was included as reference. (A) First and second principal components (dim1 and dim2) from the PCoA using all accessions analyzed in this study. (B) First and third components (dim1 and dim3) from the same PCoA. C) First and second components (dim1 and dim2) from PCoA using only S. lycopersicum var. lycopersicum samples. D) First and third components (dim1 and dim3) from the previous PCoA The percentage of explained variance for each principal component is indicated on each axis.

**Fig. 2.**
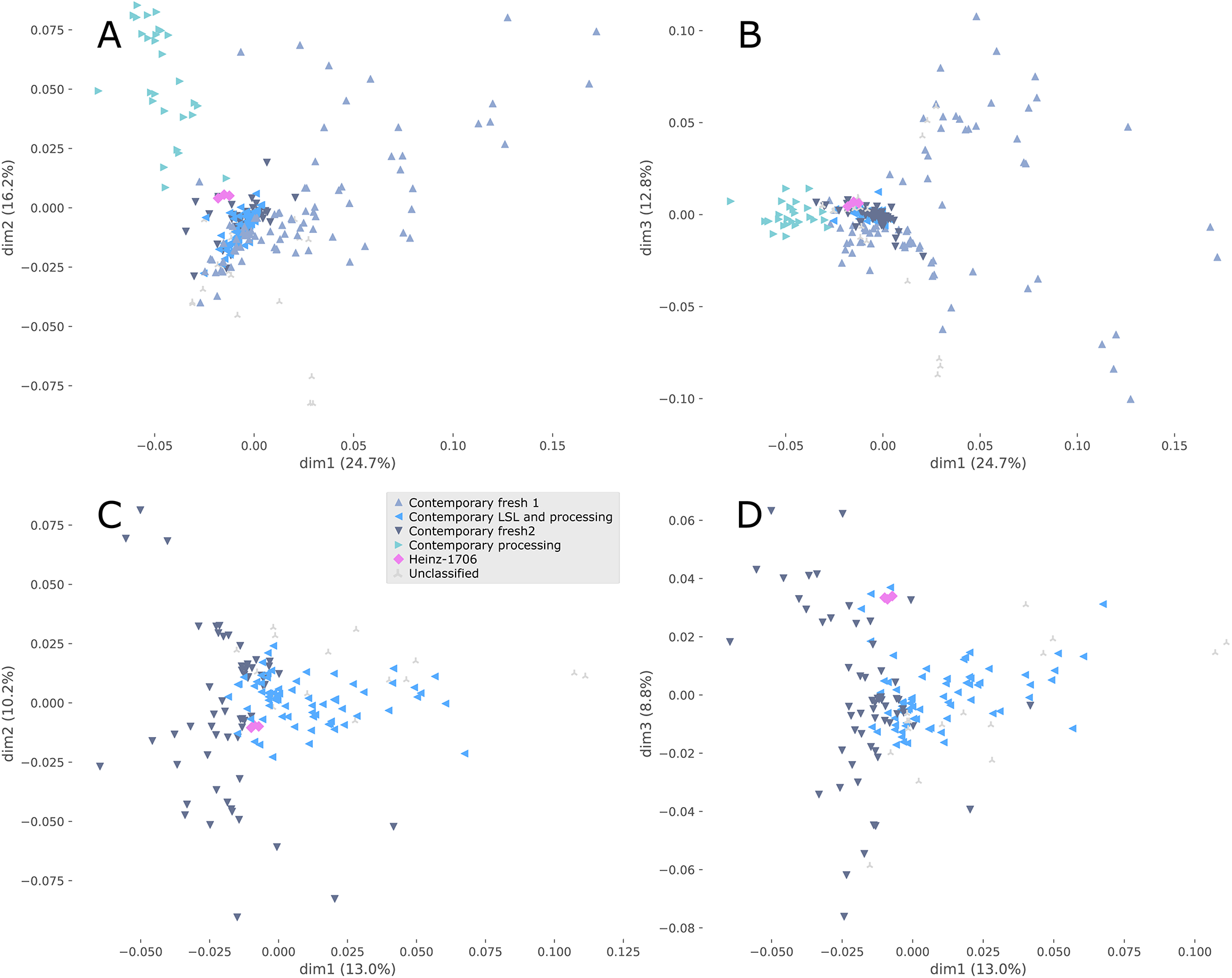
Principal Coordinate Analysis (PCoA) of modern cultivars. (A) and (B) the three first principal components (dim1, dim2 and dim3) from the PCoA considering all modern cultivars and cv. Heinz1706 as reference. (C) and (D) PCoA including only modern fresh 2 and Long Shelf Llife (LSL) and modern processing genetic groups. The variance accounted for each principal component is depicted on each axis.

The PCoA performed with this expanded collection (Fig. 1A and 1B), showed that the green fruited and Galapagos wild species, Peruvian SP (SP), Northern Ecuadorian SP (SP_NEcu), Ecuadorian SLC (SLC_Ecu), Peruvian and Mesoamerican SLC (SLC_Peru_MA), as well as several SP x SL hybrids and admixtures (SPxSL), formed a series of clusters that were clearly separated from the modern and European vintage tomatoes (Fig. 1A and 1B), with the Peruvian and Mesoamerican SLC (SLC_Peru_MA) being the closest American group to the European vintage tomatoes. To obtain a further insight into the genetic architecture of the European tomato, the genetic data was analyzed by using fastSTRUCTURE (Raj *et al*., 2014). The model marginal likelihoods reached a plateau by four populations (Supplementary Fig. 2). When this result was compared with the PCoA classification, the four fastSTRUCTURE populations were found to correspond to: SP, modern tomatoes, and two distinct vintage populations (Supplementary Fig. 2). It is remarkable that, according to fastSTRUCTURE, the modern tomato, that has been obtained after crossing varieties from different sources, was identified as an original population whereas all the wild and semi-domesticated SLCs, including the Ecuadorian, the Peruvian, and the Mesoamerican ones, appeared as admixtures.

A continuous gradient from vintage to modern rather than clearly split groups was observed in the PCoA plots (Fig. 1A, 1C and 1D). To define the limits between modern and vintage in the PCoA, we chose Heinz 1706 as the reference (in pink, Fig. 1 and 2), since it was one of the first tomato varieties reported to include introgressions from wild *Solanum* species on chromosomes 4, 9, 11 and 13 (Sato *et al*., 2012; Causse *et al*., 2013; Menda *et al*., 2014), typical of modern cultivars carrying mainly disease resistance genes.

PCoA-based classifications indicate that a total of 24.9% of the accessions labelled as vintage according to their passport data mapped outside the vintage genetic cluster in the PCoA space and were localized within the modern and SPxSL genetic groups (Fig. 1). This indicates that either they have been misclassified or correspond to a mixture between vintage and modern varieties. To find introgressions in European tomatoes, a haplotype analysis was performed to reveal haplotypes not typically found in the vintage materials. For this, the genome was divided into windows and, in every one of them, the Kosman distances were calculated from the non-vintage samples to the haplotypes found in the vintage samples. When the analyzed non-vintage sample haplotype had a non-zero distance to any of the vintage ones, it was marked as distant from the vintage collection. Several accessions mapping close to the modern varieties in the PCoA space were consistently found to include haplotypes not present in the vintage group (Supplementary Fig. 3) and, despite these being initially catalogued as vintage, it was clear that they actually came from modern breeding programs or were the result of a cross with modern cultivars, and thus were reclassified as modern genetic material (see Supplementary Table S1).

The modern materials (including both modern references and the vintage reclassified as modern) were spread across the PCoAs according to their use: fresh or processing, and also to their degree of introgression (Fig. 1C, 1D, and 2, Supplementary Fig. 3). PCoAs, when applied only to the modern accessions resulted in four groups (Fig. 2): modern processing, modern and processing long-shelf-life (LSL), modern fresh 1 and modern fresh 2 (Fig. 1 and 2). Modern processing tomatoes, the most distant group to Heinz 1706, were characterized by introgressions that included almost the entire chromosome 5 and the beginning of chromosome 11, and small introgressions in chromosomes 2, 3, 4 and 11 (Supplementary Fig. 3). Modern fresh 1 tomatoes, distributed across the PCoAs between Modern processing and Heinz 1706, were characterized by having a large introgression at the beginning of chromosome 11, a small one at the end of the same chromosome, and another introgression at the beginning of chromosome 6. Modern fresh 2 group, which is closer to Heinz 1706, was characterized by having an introgression at the beginning of chromosome 11 (Supplementary Fig. 3). The modern LSL and processing group was genetically very close to Heinz 1706 (in blue, Fig. 1C and 1D, sharing a large part of chromosome 9, including an introgression considered to be the result of the introduction of the *Tm-2* gene, conferring resistance to Tomato Mosaic Virus, in modern breeding programs. All of these haplotypes could be used for the identification of non-true European vintage tomatoes.

### Diversity among European vintage tomatoes

European vintage tomatoes are usually considered to have low genetic diversity (Blanca *et al*., 2015). Therefore, it was important to calculate the number of polymorphic variants present in our collection of European vintage tomatoes, the largest collection analyzed by sequencing thus far, and to compare it with the variability present in the wild SP, the wild and semi-domesticated SLC, and the modern cultivars. The number of variants within the European vintage collection was quite large (26,129), it was even larger than the number found in SP (19,164), in SLC (7,690), or in the materials classified as modern (17,328). However, this comparison could be biased in favor of the vintage collection because of the larger number of samples in vintage 890, compared to SP 24, SLC 42, and modern 243.

To correct for this factor, diversity indexes were calculated with the same number of samples (20) (fig 3A) and the analysis was repeated 100 times, with a different set of 20 samples chosen at random each time. Both the Nei diversity and the percentage of polymorphic variants (with a 95% threshold) was much higher in the wild SP that in any other group, and, even more relevant, both indexes were the lowest, by far, in vintage. The analysis indicated that the many of the variants found in the vintage collection could not be considered polymorphic. At 95 % threshold, the vintage collection contained only 298 polymorphic variants. This scarcity in polymorphic variants in the European vintage group can also be observed in the allele frequency spectrum (Supplementary Fig. 4) in which it is clear that most variants were almost fixed in the vintage collection.

**Fig. 3.**
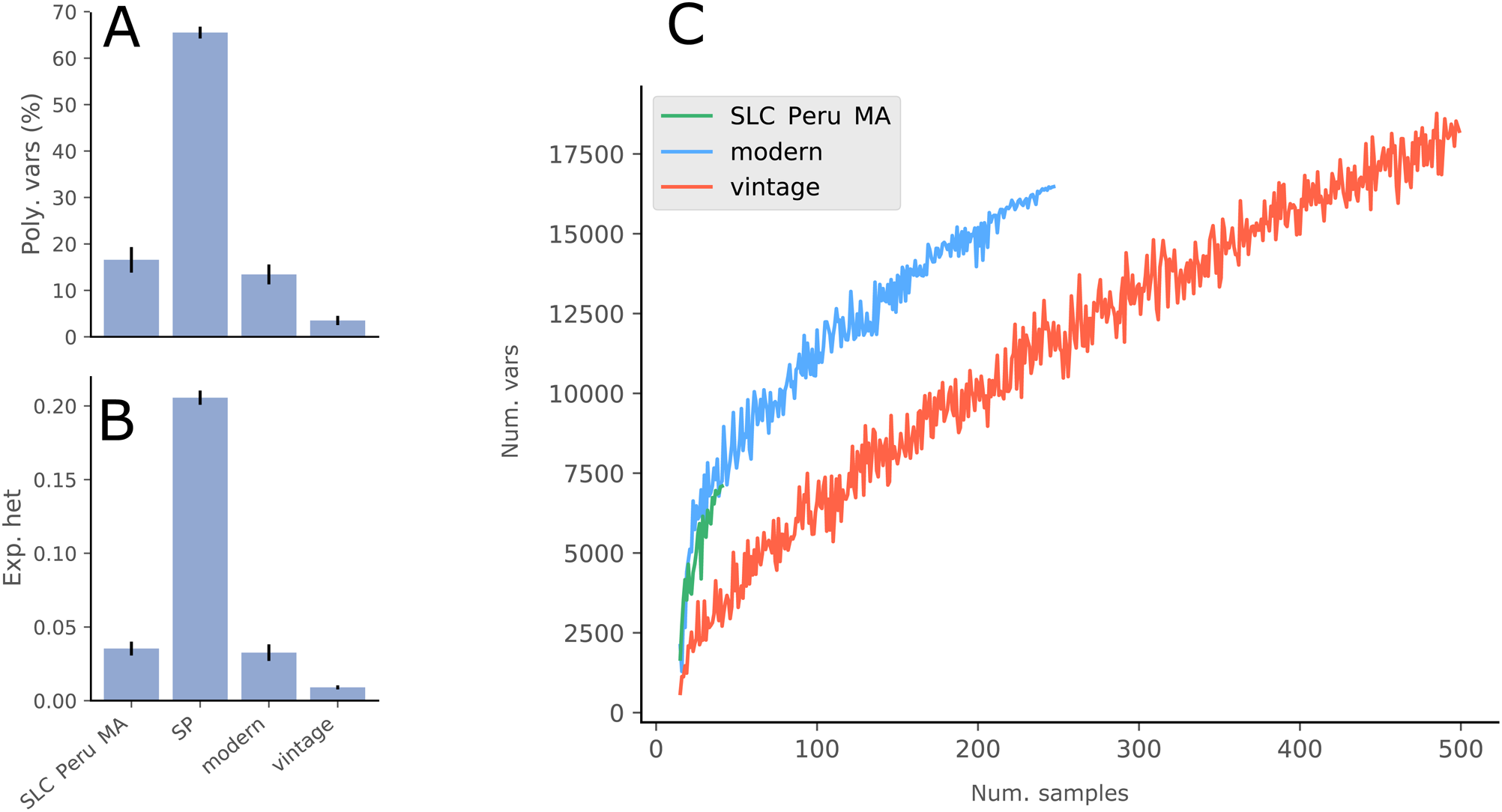
Genetic diversity for the rank1 genetic groups. (A) Genetic diversity estimated by the expected heterozygosity and the percentage of polymorphic variants (95% threshold). The indexes were calculated 100 times taking 20 samples at random from each genetic group. The mean and standard deviation are shown. (B) Rarefaction analysis of the number of variants found in each genetic group. Axis X shows the number of samples, Axis Y shows the number of variants.

To better compare the amount of genetic variability in each major cultivated group (SLC_Peru_MA, vintage, and modern) a rarefaction analysis was carried out. In this analysis, the samples were added one at a time, to check if the number of variants, including the ones at very low frequencies, reached a maximum when more samples were considered (fig 3B). The number of variants found in the vintage group was always lower than in the modern and SLC_Peru_MA groups. However, the total number of variants within the vintage collection kept increasing as more samples were added. However, the number of polymorphic variants did stabilize with a few samples. Finally, the Nei diversity decreased (Supplementary Fig. 6) when more samples were added. This decrease was due to the high number of variants found within vintage that were close to fixation.

### Linkage disequilibrium

The linkage disequilibrium (LD) was calculated for the genetic groups with enough polymorphic markers (Minimum Allele Frequency (MAF)> 0.02 threshold) (Supplementary Fig. 6), which were SP, SLC from Peru and Mesoamerica, modern, and European vintage varieties. Wild SP showed the lowest LD (r^2^=0.42) at 5 kb and it was also the group in which LD decreased the fastest, being only r^2^=0.2 at 25 Kb. In SLC, r^2^ was 0.8 at 5 kb and at 1000 Kb it was still 0.4. Vintage had the highest LD at 25 Kb (r^2^=0.97); however, it decreased to the lowest value (r^2^=0.05) at 1000 Kb. The modern accessions had a high LD both at 25 Kb (r^2^=0.9) and at 1000 Kb (r^2^=0.6). The LD found at 1000 Kb is likely due to population substructure. SLC and modern had high long range LDs, perhaps because modern included both fresh and processing accessions, which were clearly separated in the PCoAs, and SLC contained accessions from Peru and Mesoamerica, two geographically distant areas. Additionally, modern cultivars often contain introgressions from wild species, including disease resistance genes, that span large regions for which recombination is usually suppressed. SP is also known to have a clear population structure (Blanca *et al*., 2012) and also showed some long range LD, which clearly supports the conclusion that LD is not due just to gamete disequilibrium, but to other causes too. The vintage accessions showed the lowest LD at 1000 Kb perhaps because it has a less remarkable population substructure.

### Classification of vintage tomato clusters

To further classify true vintage tomatoes, a series of PCoAs (Supplementary Fig. 7) were performed. A genetic group was created when several samples that grouped together in the PCoAs shared their geographic origin or traditional variety name, or some aspect of their phenotype (Supplementary Table S2) e.g. fruit shape and size. Most vintage samples could be classified into 27 different genetic groups, using this PCoA strategy, and were named as *“Balearic cherry”*, “*Bell pepper”, “Cor de bou”, “Greek (grc)”, “Italian (ita) ellipsoid”, “Ita grc”, “Ita small”, “Lemonia”, “Liguria”, “Long Shelf Life (LSL) da serbo”, “LSL heart”, “LSL penjar cat”, “LSL penjar vlc”, “LSL piennolo”, “LSL ramellet”, “Marmande”, “Montserrat”, “Muchamiel”, “Palosanto pometa 1”, “Palosanto pometa 2”, “Pera girona”, “Pimiento”, “San Marzano”, “Scatolone di bolsena”, “Spagnoletta”, “Tondo piccolo”, “Valenciano”*.

Two connected clusters of genetic groups (for the sake of clarity we will use “group” to refer to a PCoA group of samples and “cluster” to talk about a cluster of groups) were observed in PCoA (Supplementary Fig. 7A and 7B). Within the cluster at the center of PCoA, we found the genetic groups *“LSL ramellet”, “LSL penjar vlc”, “LSL penjar cat”, “LSL ramellet”, “Marmande”, “Montserrat”*, “*Bell pepper”, “Lemonia”, “Muchamiel”, “Palosanto pometa 1”, “Palosanto pometa 2”, “Pera girona”, “Scatolone di bolsena”, “Spagnoletta” and “Valenciano”* (Supplementary Fig. 7C and 7H). These genetic groups belong to Spain, with the exception of *“Marmande”*, “*Bell pepper”* and *“Palosanto pometa 1”*, which were represented in all four Mediterranean countries (Spain, Italy, France and Greece), the Italian *“Scatolone di bolsena”* and *“Spagnoletta”*, and the Greek “*Lemonia”* (Fig. 4A). Outside the central cluster, but close, we found groups of big tomatoes: “*Liguria*”, with accessions mainly collected in Italy, and “*Cor de bou*” and “*Pimiento*”, present in all four countries (Supplementary Fig. 7C and 7D, Fig. 4A and 4B). A second cluster included mostly Italian accessions classified into the *“Ita ellipsoid”, “Ita small”, “LSL da serbo”, “LSL piennolo”, “San Marzano”*, and *“Tondo piccolo*” genetic groups, and also some Greek and Spanish accessions included in the *“grc”, “Ita grc”, and “Balearic cherry”* genetic groups, all characterized by having a small size with no or weak ribbing (Supplementary Fig. 7A, 7B. 7M-R, Fig. 4A and 4B). In summary, the PCA separated vintage accessions mainly by country of origin and fruit size. It is interesting to note that the LSL-type accessions, which were highly represented in the collection, were not grouped together, but rather segregated by country: the Italian LSL varieties were found within the Italian cluster, and the Spanish LSL within the Spanish cluster. Several accessions located between the Spanish and the Italian clusters could not be grouped by passport data or any other characteristic.

**Fig. 4.**
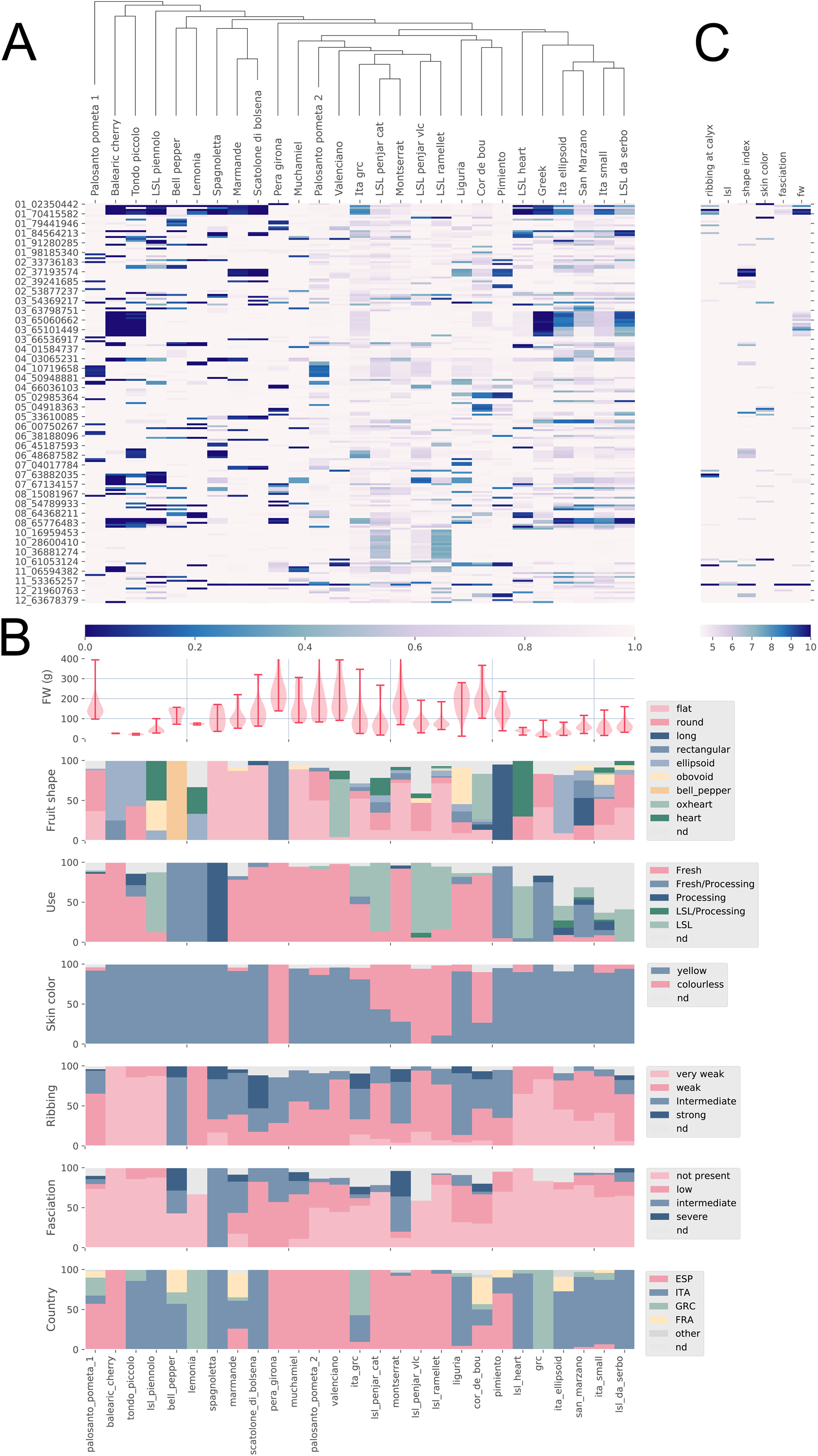
Allele frequencies across the genome in Vintage genetic groups and their relationship with phenotypic diversity. (A) Clustering of genetic groups based on allele frequencies. Allele frequency of the major allele within each genetic group is indicated by a density color according the legend (blue, frequency=0, to white, frequency=1. (B) Distribution of the different traits within genetic groups. (C) Statistical significance indicated by a colored gradient of -log(p) values of the SNP-trait associations by Genome-Wide Association Analysis.

### Allele frequencies across the genome in Vintage groups and their relationship with phenotypic diversity

A clustering of the vintage genetic in groups based on a distance tree was calculated using the polymorphic variants (95% threshold) (Fig. 4A). This analysis showed that the defined genetic groups had quite distinct allele frequencies along the genome. Concomitantly, the genetic groups also showed enrichment in specific phenotypic characteristics related to their horticultural classification. For example, varieties belonging to the genetic groups “*Pera girona*”, “*LSL ramelle*t” and “*LSL penjar vlc*” have colourless skin, while “*Balearic cherry*”, “*Tondo piccolo*”, *LSL piennolo*”, “*Lemonia*”, “*LSL heart*”, “*LSL Penjar vlc*”, “*grc*”, and “*San Marzano*” showed mostly weak ribbing, and, finally, “*Spanoletta*” was characterized by its fasciation (Fig. 4B). Moreover, the fruit size was also different for different genetic groups and clusters *“LSL heart”*, “*Ita ellipsoid”, “Ita small”, “LSL da serbo”, “LSL piennolo”, “San Marzano”, “Tondo piccolo*”, *“grc”, “Ita grc” and “Balearic cherry”* were characterized by having a small size, while the rest were medium or large in size. Furthermore, several noticeable clusters of genetic groups with common phenotypic traits could be observed. For instance, there was a cluster formed by small-fruited, slightly-ribbed, long shelf-life and processing Italian genetic groups which included the well-known Italian *“da Serbo”* and *“San Marzano”* tomatoes. Another cluster was comprised mainly by long shelf-life colourless-skinned Spanish tomatoes, which included the *“LSL penjar cat”, “LSL penjar vlc”*, and the *“LSL ramellet”* groups. Interestingly, this cluster also included the Catalonian big fruited *“Monserrat”* group which, in contrast to the others, were fasciated and used for fresh consumption. Close to this cluster were some of the most typical Spanish vintage fresh-market varieties: *“Valenciano”, “Muchamiel”* and *“Palosanto pometa 2”*. In addition, big tomatoes appertaining to *“Liguria”*, “*Cor de bou*”, and “*Pimiento*” clustered together.

Some of the genetic differences between the groups could be due to genetic drift not related with the phenotypic variability generated during the history that gave rise to the different vintage varieties, but allele frequencies of genes involved in the phenotypic variation could have been selected either inadvertently or consciously by traditional farmers. In order to elucidate whether the differentiating variants were associated to the phenotypic variation observed in the different genetic groups a GWAS analysis was carried out using selected fruit characters (Fig. 4C). Two of the main phenotypic characteristics differentiating the vintage tomatoes are fruit weight (fw) and ribbing (Fig. 4B). In the GWAS analysis fruit weight was associated to variants on chromosome 1, 3 and 11. The MAF analysis indicated that most of the small fruited tomatoes such *“LS heart”*, “*Ita ellipsoid”, “Ita small”, “LSL da serbo”, “LSL piennolo”, “San Marzano”, “Tondo piccolo*”, *“grc”, “Ita grc”*, and *“Balearic cherry”* shared the fixation of the same allelic variant in chromosome 1. The pattern found in chromosome 3 was similar, except for the *“LS heart”* and “*LSL piennolo”* groups.

For ribbing, GWAS revealed association with variants on chromosomes 1, 7, 10 and 11. The chromosome 1 region was fixed in the weak ribbed groups *“Balearic cherry”, “Tondo piccolo*” and *“LSL piennolo”*. In contrast, almost all medium and large tomatoes, with the exception of *“Pimiento”* and “*Spanish LSL*” fruits (both showing no or weak ribbing) had a fixed common variant in chromosome 11 that was associated by GWAS with fruit weight, ribbing at calyx end, and fruit shape index.

Another trait differentiating vintage tomato cultivars was skin colour, for instance, most Spanish LSL as well as tomatoes included in the *“Cor de bou*”, *“Montserrat”*, and “*Pera girona*” genetic groups had a colourless skin, which resulted in pinkish fruit (Fig. 4B). GWAS found association with this pink color in chromosomes 1 (two regions), 3, 5, and 10. The GWAS and MAF analysis comparison (Fig. 4A and 4C) showed that different pink genetic groups had different allelic composition in the associated genomic regions, what might reflect a complex genetic control. Fruit shape was associated with regions in chromosomes 2, 5, 10, and 12. The region in chromosome 2 was fixed in *“Marmande*” and “*Scatolone di bolsena*”, two groups that are well known for having flat fruits. In addition, “*Valenciano*”, “*Pimiento*”, and “*Liguria*” had the minor allele almost fixed in the chromosome 10 region. High frequency minor alleles, almost fixed in the regions associated to fruit shape in the GWAS, were also observed in other genetics groups such as the Italian “*LSL da serbo*”, in chromosome 5, and “*Ita ellipsoid*” and “*Tondo Piccolo*”, in chromosome 6 as well as in “*Cor de bou*” and *“Pimiento*” groups, in chromosome 12.

In the case of use, associated variants were found in chromosomes 10 and 11, but, in this case, no clear relationship was found between allelic frequencies among the tomato genetic groups and GWAS.

### Network analyses supports the differentiation between Spanish and Italian vintage tomatoes and the occurrence of hybridization events in vintage tomatoes across Europe

To study the genetic relationships between accessions and groups of accessions, a network based on pairwise Dest group distances was created with Splitstrees. Evolutionary relationships are often represented as an unique tree under the assumption that evolution is a branching or tree-like process (Husson 1998). However, real data does not always clearly support a tree. Phylogenetic split decompositions represented in a network may be evidence for conflicting reticulated phylogenies due to gene flow and/or hybridization (Husson 1998).

The splitrees network of European tomato is depicted in Fig. 5. The group organization in the network was structured, like the PCoAs (Supplementary figure 7), in two main country-related clusters. One cluster was comprised mainly of Spanish vintage groups, which included the Spanish LSL, *“Muchamiel*”, and *“Montserrat”* types, and another cluster was mostly comprised by the small fruited Italian LSL and processing groups. Interestingly, the “*Liguria*” group clustered with Spanish clusters, although the branch that linked it with the core Spanish clusters was quite large.

**Fig. 5.**
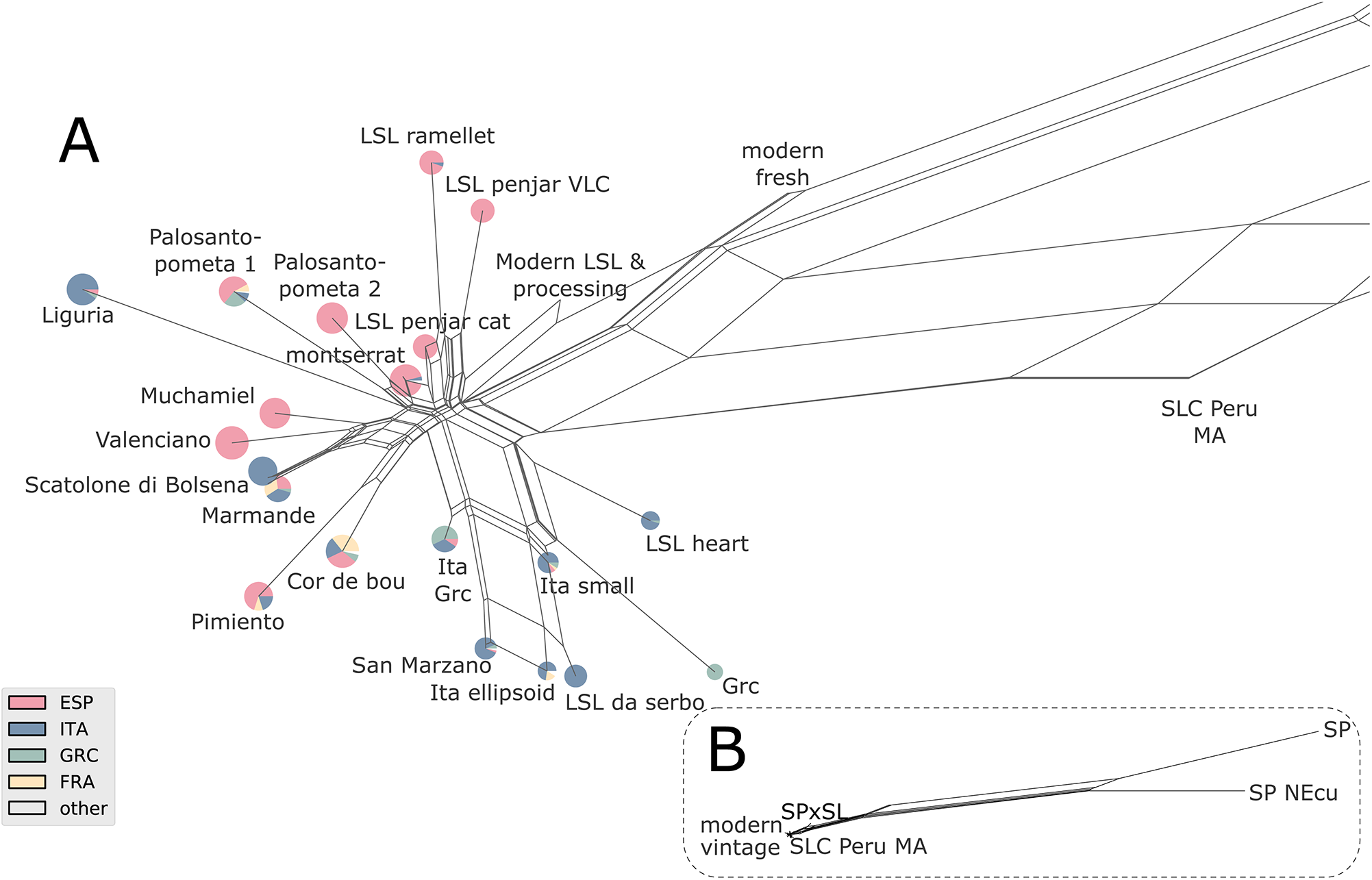
Evolutionary relationships between vintage European tomato, Modern tomato and Peru and Mesoamerica *Solanum lycopersicum* var. *cerasiforme* (SLC), *S. pimpinellifolium* (SP) and the hybrids SPxSL. Split network based on the Dest distances between genetic groups. The country of origin of accessions within each genetic group is represented by a pie chart depicted in the bottom left. (A) Zoom only in European modern and vintage tomatoes. (B) Zoom on American ancestral and wild tomatoes. Each edge of the network represents a split of the accessions based on one or more characteristics. If there was no conflict, each split was represented by a single edge, while in the case of contradictory patterns the partition was represented by a set of parallel edges. The edge lengths represent the weight of each split, which is equivalent to the distance between groups.

The degree of reticulation found (Fig. 5) suggested that hybridizations might have occurred between the ancestors of accessions collected from the same geographical regions. On the other hand, the groups that included accessions from different countries, such as “*Marmande*”, “*Pimiento*”, “*Cor de bou*” or “*Palosanto pometa 1*”, were located between the Spanish and Italian clusters.

These groups of mixed origin could be more modern and derived from hybridization from old Spanish and Italian varieties or, alternatively, they could be very old varieties found across Europe before the Spanish and the Italian diversification started. To check those possibilities, a rarefaction analysis was performed of the number of polymorphic sites found in these three clusters was calculated (Supplementary Fig. 8). The number of polymorphic sites was clearly higher in the Italian and Spanish clusters and much lower in the mixed origin cluster, an evidence that supports that Spain and Italy were secondary centers of diversity for the European tomato, whereas the varieties included in the mixture cluster would be more recent.

## Discussion

### Very low, but discriminant, variation in vintage European tomatoes

The genetic diversity of this European vintage collection was very low when compared with the diversity found in SP or even in SLC. While this result is in agreement with previous surveys on worldwide SLL accessions carried out with the SolCAP SNP platform (Sim *et al*., 2012, Blanca *et* al., 2012; Blanca *et al*., 2015), the current analysis represents the first estimate obtained using a comprehensive representation of vintage European tomatoes, and it is relevant to study the role of Europe as a secondary center for tomato diversification. The low level of diversity found in these traditional materials was quite striking: after sequencing 0.8% of the genome, only 298 polymorphic variants at the 95% level were found. This result is quite remarkable when we consider the high phenotypic diversity of vintage tomatoes. Moreover, the high linkage disequilibrium found in these traditional vintage materials suggests that it is rather unlikely that the total number of polymorphic blocks would grow much even if whole genome sequences were to be obtained.

Previous studies demonstrated a strong bottleneck during the SLC tomato’s travel from Ecuador and Peru to Mesoamerica (Blanca *et al*., 2015, Lin *et al*., 2014; Razifard *et al*., 2020). However, it is remarkable that despite the low genetic diversity found in vintage European tomatoes there are still a few highly polymorphic loci within this tomato gene pool. Some of this variation could be due to the random nature of genetic drift. However, the association study carried out with major phenotypical/morphological traits found that a sizeable fraction of those diverse loci were associated with the vintage fruit phenotypical/morphological variation. Therefore, it is quite likely that many of those polymorphic loci had been under balancing selection (Delph and Kelly, 2014) during the diversification process and were in fact responsible for a sizeable part of the tomato phenotypic variation, or, at least, in linkage disequilibrium with the variants selected. It may seem paradoxical that the high diversity of shapes, colors, sizes, uses, and other agronomic traits in the vintage group could be maintained by such a poor gene pool, but it seems that the selection carried out by the traditional growers in favor of this agronomic diversity resulted in a desert of variation, with just a handful of scattered polymorphic loci. This is consistent with two highly polymorphic SNPs found by Muños *et al*., (2011) *in the lc* locus. These were highly polymorphic, but were surrounded by loci with “drastically reduced” diversity. Thus, they seemed to be the result of selection for low or high number of locules in different materials.

Recently, structural variants (SV) were studied in tomato using new long-read sequencing technologies and new analysis algorithms (Alonge *et al*., 2020; Domínguez *et al*.,, 2020). A large number of structural variants were identified and were mostly generated by transposons and related repeats. Similar to the variants studied here, most structural variants had a very low frequency, and the majority were singletons.

Therefore, the phenotypic diversity present in European vintage tomatoes seems to have been built by remixing/reshuffling/swapping very few polymorphisms with the selection pressure associated with the creation of new varietal types and to the adaptation of these types to different regional environments.

### Tomato History: tomato movement in Europe

The distribution of the genetic variability in the European vintage tomatoes showed mostly a continuous gradient. However, the Spanish and Italian varieties occupied opposite regions of the PCoA space what supports a genetically differentiation among varieties originated in those countries. The lack of clear-cut limits may be due to migrations between different regions and countries and subsequent intercrossing. Despite this difficulty, the genetic vintage groups proposed here were differentiated by characteristics such as: their main geographic origin, use, fruit morphology, and varietal name. The genetic groups sometimes corresponded with the varietal type, such as in “Valenciano”, “Muchamiel”, “Penjar” or “Piennolo”. However, the match between the proposed genetic group and the sample varietal name was seldom complete. For instance, the “*Cor de bou*” group included two “Valenciano” samples, one “Russe”, and one “Costoluto”. This may be due to the limitations of the genotyping or genetic classification methodology utilized or to erroneous passport data, as it may not be trivial for a standard grower to evaluate the sometimes subtle varietal differences. Other genetic groups, such “Italian small” showed no clear associations to any variety name.. Finally, cultivars previously classified as belonging to some variety, such as “Marmande”, were included in many different genetic groups. It is likely that the popularity of some varietal types such as “Marmande”, made some growers prone to apply the label to any variety that displayed the typical morphological characteristics of a well-known varietal type. Thus, the “Marmande” tomatoes are characterized by its production of large and multi-locule tomatoes, and any other variety with a similar fruit morphology could have been labeled as “Marmande”.

One clear example of mistaken identities and/or inadvertent out crossing is provided by the vintage samples that were found to include haplotypes not found in the vintage core and to be genetically closer to the modern varieties than to the vintage materials in the PCoA. It is not even trivial to define the borderline between vintage and modern varieties. One could think that until the 1950’s most varieties were heirlooms and landraces maintained by small farmers, but the real history is more complex. When tomato cultivation was popularized in the 19th Century in France, England, and the USA some of the varieties were already provided by seed companies (Boswell 1937), and there were seed shipments documented between countries, for instance, from England to the Canary Islands (Amador *et al*., 2012). Moreover, from 1910 onwards, professional breeding efforts created new varieties adapted to long-distance shipping and for processing (Boswell 1937). These efforts did not yet include wild materials, so their results are not easy to be differentiated in a PCoA analysis. It is only when shortly afterwards, breeders started introgressing wild species alleles for disease resistance, that the varieties created were different enough to be easily differentiated in the PCoA analysis. In any case, the vintage-modern limit has to be somewhat conventional, although a characteristic of modern cultivars compared with vintage varieties is the introgression of genes from wild species. Therefore, true vintage cultivars were defined based on the absence of wild species haplotypes.

Most of these introgressions seem to be related to disease resistance genes as the *Cladosporium fulvum* resistance gene *Cf-2* in chromosome 6, *Tm-2* (resistance Tomato Mosaic virus,) in chromosome 9. It is likely that the modern genetic variability has been combined with the true traditional varieties, so some materials catalogued in the genebanks as traditional are in fact a mixture of traditional and modern. This is to be expected, as the seed collectors/genebanks label as vintage any material considered as such by the farmer from whom the seeds where collected. Although European small farmers often save their own tomato seed, they may occasionally purchase or get plantlets from markets or nurseries or save seeds from modern varieties purchased in the market and introduce them in their fields. This may lead after several years of reproduction and farmer selection to complex hybridizations and mixings. Clearly, there have been many opportunities for introgressing modern haplotypes into the vintage materials, such as unintentional crosses. This phenomenon could be thought of as blurring the boundaries of a supposedly pure vintage population, but one may also think that this leakage had the positive unintended consequence of increasing the very low diversity of the vintage pool, and it is also the case that evolution consists of change and adaptation of local varieties (Casañas *et al*.,, 2017).

The allele frequency based tree (Fig. 4) defined three major clusters: Spanish, Italian, and Mixed origin. The mixed origin groups are basal in the Fig.4 tree, have longer branch lengths, and occupy an intermediate position between the Italian and Spanish clusters in the Dest network (Fig. 5). These results are compatible with the hypothesis that Italy and Spain formed two centers of diversity. The differentiation of Italian and Spanish gene pools is exemplified by the long LSL varieties from both countries. Italian and Spanish LSL varieties were clustered apart from each other with only a small number of samples from the other country, so it seems as if the origin of the long shelf life tomatoes in both countries was independent. The transformation from a fresh to a long shelf-life variety is likely due to a limited number of loci, as observed in Fig. 4 in which the Catalonian fresh “Montserrat” type is closely related to the Catalonian long shelf-life “Penjar” type. Esposito et al (2020) also observed geographic differentiation of the Italian and Spanish long shelf-life varieties. Therefore, although there may have been migrations from Italy to Spain and vice versa, these may not have been extensive enough to erase the genetic differences between the Italian and Spanish varieties

Regarding the mixture cluster, the groups included in it are basal in the Fig.4 tree, they have longer branch lengths, and occupy an intermediate position between the Italian and Spanish clusters in the Dest network (Fig. 5). Moreover, the rarefraction analysis supports that it included varieties derived from the two secondary centers of diversity. This could be the result of long of tomato cultivation tradition in Southern Europe, being the groups included in this cluster developed from hybridizations between the two centers of diversity. New mutations, other introductions of tomatoes from America or new genes from varieties developed worldwide might also be involved in the history of the groups of mixed origin.

A complex pattern of migrations can also be inferred in several genetic groups as the “*Cor de bou*” group that included varieties from most countries: French “Coeur de boeuf”, Italian “Cuor di bue”, Catalonian “Cor de bou”, Castillian “Corazon de toro”, and “Navarran corazón de fitero”. Also, the Italian “Spagnoletta” group was closely related with the “*Marmande*” group comprised by French, Spanish, Greek, and Italian accessions. Other genetic groups with mixed geographic origin are “*Liguria*”, “*Cor de bou*”, “*Pimiento*”, “*Palosanto Pometa 1*”and “*Marmande*”.

### Do a few Polymorphic genes differentiate the true European vintage tomato genetic groups?

In order to shed light on the apparent contradiction between the low genetic diversity and the large phenotypic variation of European vintage tomatoes, a GWAS was carried out with the polymorphic variants and some of the most obvious morphological traits (fruit morphology, color, and ripening behavior).

Variants located in the genomic regions of previously identified loci involved in fruit weight, and likely involved in domestication and diversification, were associated with this trait in the GWAS performed with the European vintage collection. Most of the small fruited tomatoes shared fixed variation regions in chromosomes 1 and 3 which mapped close to previously-described Quantitative Trait Loci (QTL)and genes associated with fruit size: *fw1*.*1* (Grandillo *et al*., 1999) and *fw3*.*2/KLUH* and *ENO* (Chakrabarti *et al*., 2013; Yuste-Lisbona *et al*., 2020) (Fig. 4A and 4B). In contrast, almost all medium and large tomatoes shared a region in chromosome 11 that mapped close to FAS (Xu *et al*., 2015) and *fw11*.*3*/CSR (Mu *et al*., 2017), with both genes playing a known role in controlling fruit size and fasciation. No more associations were observed in other genomic regions for fruit weight, so it seems reasonable to think that these QTLs can be responsible, at least in part, for the fruit size variability among the European vintage tomatoes. Regarding fruit shape, two of the associated regions found included known genes. The chromosome 2 region included previously mapped QTLs as heart shape *hrt2*.*2* (heart shape), *pblk2*.*2* (proximal end blockiness), *psh2*.*2* (shoulder height), *piar2*.*2* (indentation area) (Brewer *et al*., 2007) and *ovate* (Liu *et al*., 2002), *and the region in chromosome 10 is located close to, where the original copy of the sun* locus was found (Xiao *et al*., 2008). In the case of skin color, a different pattern was characteristic of different pink genetic groups. “*LSL Penjar* vlc” and “*LSL ramellet*” shared a variant at the end of chromosome 1 that matched a region that was previously associated with skin color, the colorless-peel and locus (Ballester *et al*., 2010), while “*Pera Girona*” had the minor allele for the other chromosome 1 variant, which is located at the beginning of the chromosome and maps close to the SlCMT3 (Gallusci *et al*., 2016) and PSY3 (Li *et al*., 2008) genes involved in epigenetic ripening regulation and carotene biosynthesis, respectively.

The current analysis suggests that fruit morphology variability among European vintage tomatoes could be the consequence of the combination of a relatively low number of genes, as suggested by Rodriguez *et al*., (2011), including *fw3*.*2/KLUH, ENO, FAS*, SUN, and OVATE. On the other hand, skin color could be a consequence of y-locus and other genes related to phenylpropanoid metabolism. Interestingly, SV mutations have been found in *fw3*.*2/KLUH, FAS* and *SUN* that supports the impact of SV on tomato phenotypic diversity (Alonge *et al*., 2020, Dominguez *et al*., 2020). Also some cryptic variation hidden in the Mesoamerican tomatoes may have emerged in European tomatoes after generating new combinations and divergent selection by the traditional farmers as found for the jointless trait in tomato (Soyk *et al*., 2017, Soyk *et al*., 2019, Alonge *et al*., 2020).

### Impact on genebank and on farm variability management

Many of the few polymorphic genetic variants, within the very low diversity European vintage tomatoes, appeared to be associated with phenotypic variation. This has implications for the conservation efforts carried out by the genebanks. Thousands of European vintage tomatoes are maintained in many of those genebanks. However, the cost of these conservation efforts could be severely reduced if only these few polymorphic loci were taken into account. Of course, that would ignore most variants, the ones found in very low frequencies, but conserving these low frequency alleles, that in many cases would be neutral, and thus not associated with any phenotypic variation, requires a sizeable investment. An alternative would be to identify the alleles associated with a phenotype, however, that would require an exhaustive phenotypic characterization.

Most of the European accessions analyzed here were collected from farmers in the 1950’s to 1980’s, and as landraces, they are appreciated, competitive, and cultivated varieties. The genetic diversity of many other crops has also been maintained as landraces that evolved on-farm. However, this diversity is continuously under threat by the introduction of new modern varieties derived from a limited gene pool that have replaced the vintage varieties. It is generally believed that most of the accessions in seed banks do not contribute to modern varieties (Tanksley and McCouch, 1997) and this is also the case for tomato. Our Identification of the morphological and genetic structure present in the European vintage tomato gene pool will be important to guarantee access to that variability as the basis of the development of new varieties or evolved landraces in the future (Casañas et al 2017).

## Conclusion

The entrepreneurship of many local European farmers during the last five hundred years has managed to create a very complex and diverse set of tomato varieties adapted to different local tastes and morphological preferences. These localized activities did not restrain those farmers from importing other interesting novelties developed by other farmers elsewhere, thus generating a much larger set of varietal tomato types that are characterized by an exuberant diversity that serves as a variety for fresh, processing, and long shelf-life uses.

The current report shows that such a plethora of different types has been created from an original material devoid of genetic diversity, by exploiting very few polymorphic loci subjected to balancing selection.

## Supplementary Data

Fig. S1 Number of genomic positions with high coverage and number of variants per megabase along the genome in all accessions.

Fig. S2. FastSTRUCTURE analysis.

Fig. S3. Introgressed regions along the genome detected in the modern genetic groups

Fig. S4. Major Allele Frequency spectrum in vintage, modern, and SCL_Peru_MA

Fig. S5. Rarefaction analysis of the expected heterozygosity for each genetic group

Fig. S6. Genome-wide linkage disequilibrium (LD) decay in wild, *S. lycopersicum* var. *cerasiforme*

(SLC), vintage, and modern accession groups.

Fig. S7. Hierarchical Principal Coordinate Analysis of European vintage tomato varieties

Fig. S8. Rarefaction analysis of the number of polymorphic variants (95% threshold)

Table S1. Accessions analyzed in this study.

Table S2. Phenotypic characterization of European vintage tomatoes

Table S3. Sequencing and mapping statistics for each sample.

## Acknowledgments

Authors want to thank Maurizio E. Picarella for expert technical assistance. CNR-IBBR acknowledges the seed donors ARCA2021, Cristian Patané (CNR-IVALSA, Catania, Italy), La Semiorto Sementi SRL and the Leibniz Institute of Plant Genetics and Crop Plant Research and the Gene Bank of DNR-IBBR (Bari, Italy). We also thank Universitat Illes Balears, the Greek Gene Bank (GGB-NAGREF), Università degli Studi Mediterranea Reggio Calabria for seed sharing. This work was supported by European Commission H2020 research and innovation program through TRADITOM grant agreement No. 634561, G2P-SOL, grant agreement No. 677379, and HARNESSTOM grant agreement No. 101000716. Mariola Plazas is grateful to Spanish Ministerio de Ciencia e Innovación for a post-doctoral grant (IJC2019-039091-I/AEI/10.13039/501100011033).

## Author contributions

JB and JC analysed the data and drafted the manuscript. J M-P, D S-M, PZ, RF analysed the data. CP obtained the DNA, field trial phenotypic data and revised manuscript.

LF, JF, MP, JLR, ARiccini, SP, ARuggiero and MS obtained field trial phenotypic data. JC obtained field trial phenotypic data, selected and provided vintage varieties.SG, AK, GG, MC, SG, AM, MC, MJD, JP, selected and provided vintage varieties and revised the manuscript. DZ coordinated the field trial. AJM and AG conceived and coordinated the study and revised the manuscript.

## Data Availability

The sequence data can be found in NCBI (https://www.ncbi.nlm.nih.gov/sra) under the accession number PRJNA722111.

## Abbreviations

GBS: Genotyping by Sequencing
GWAS: Genome-Wide Association Analysis
LD: Linkage Disequilibrium
LSL: Long Shelf-Life
MAF: Minimum Allele Frequency
PcoA: Principal Coordinate Analyses
QTL: Quantitative Trait Locus
SLL: *Solanum lycopersicum* L. var. *lycopersicum*
SLC: *S. lycopersicum* var. *cerasiforme*
SNP: Single Nucleotide Polymorphism
SP: *S. pimpinellifolium*

