## Supplementary material for "European Vintage tomatoes galore: a result of farmers combinatorial assorting/swapping of a few diversity rich loci": Supplemetal figures

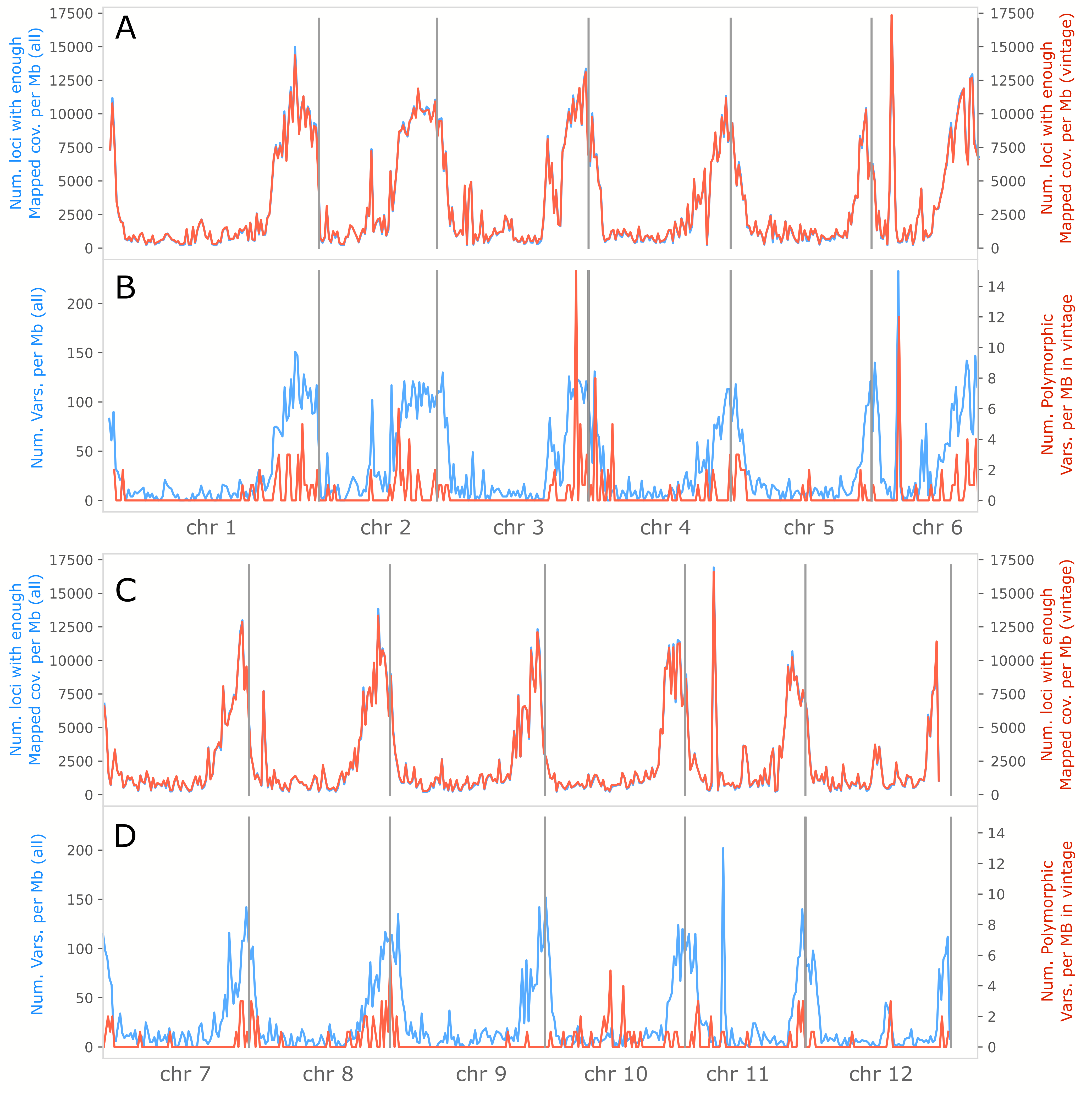
A
B
C
D
Num. loci with enough
Mapped cov. per Mb (all)
Num. loci with enough
Mapped cov. per Mb (vintage)
Num. Vars. per Mb (all)
Num. Polymorphic
Vars. per MB in vintage
chr 1
chr 2
chr 3
chr 4
chr 5
chr 6
chr 7
chr 8
chr 9
chr 10
chr 11
chr 12

Fig. S1. Number of genomic positions with high coverage and number of variants per megabase along the genome in all accessions used in this study (blue line) and in vintage European tomato (red line). (A, C) Number of loci with enough reads mapped per Mb. (B, D) Number of polymorphic sites per Mb.

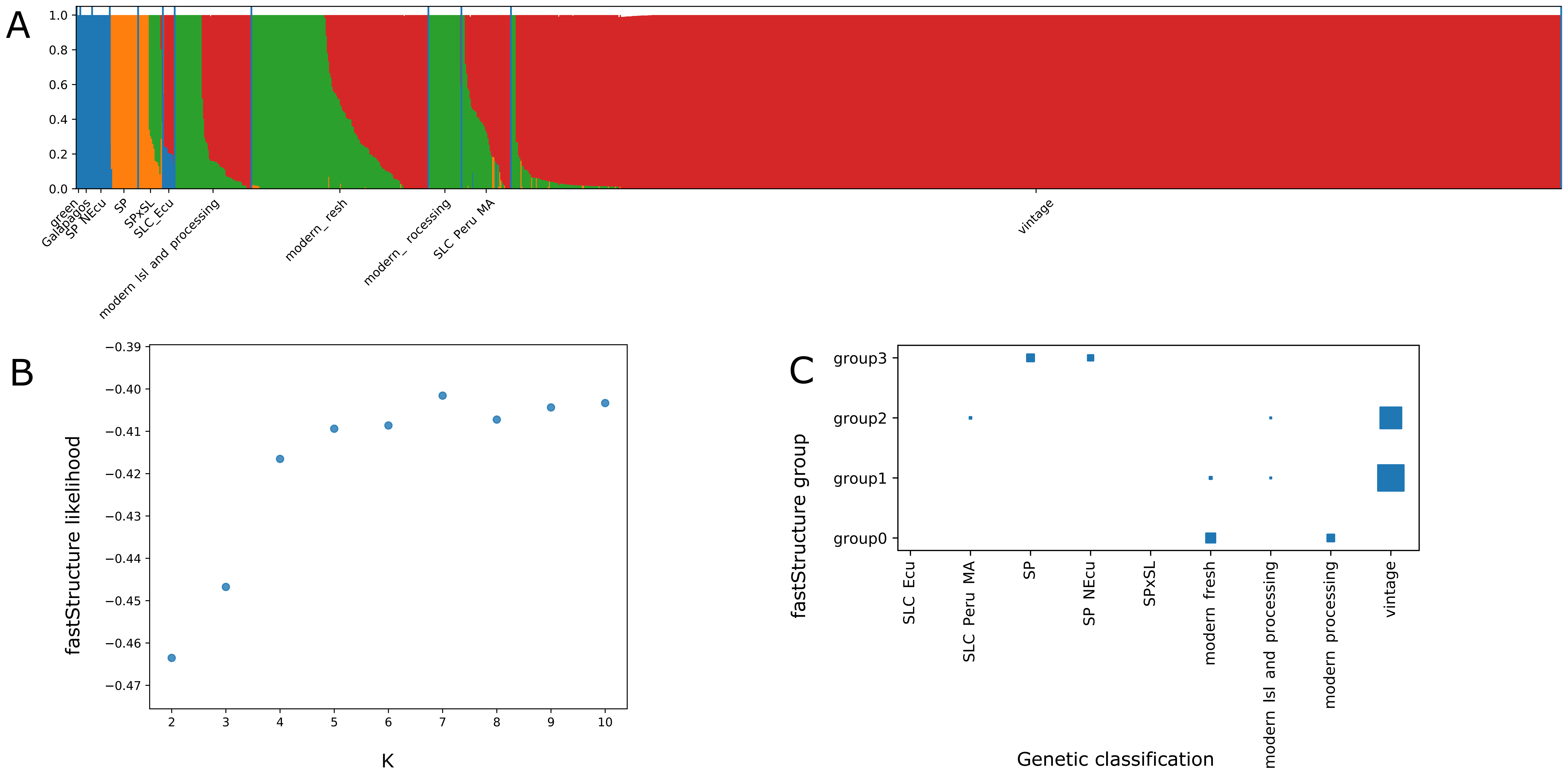
A
1.0
0.8
0.6
0.4
0.2
0.0
green
Galapagos
SP NEcu
SP
SPxSL
SLC_Ecu
modern Isl and processing
modern_ resh
modern_ rocessing
SLC Peru MA
vintage
B
fastStructure likelihood
−0.39
−0.40
−0.41
−0.42
−0.43
−0.44
−0.45
−0.46
−0.47
2
3
4
5
6
7
8
9
10
K
C
fastStructure group
group3
group2
group1
group0
SLC Ecu
SLC Peru MA
SP
SP NEcu
SPxSL
modern fresh
modern Isl and processing
modern processing
vintage
Genetic classification

Fig. S2. FastSTRUCTURE analysis. (A) Assignation of the accessions to faststructure groups for K=4 according with Structure likelihood saturation for different Ks (B). (C). Comparison of the Structure classification (k=4) and the PcoA-based classification. The number of samples is represented by the square surface.

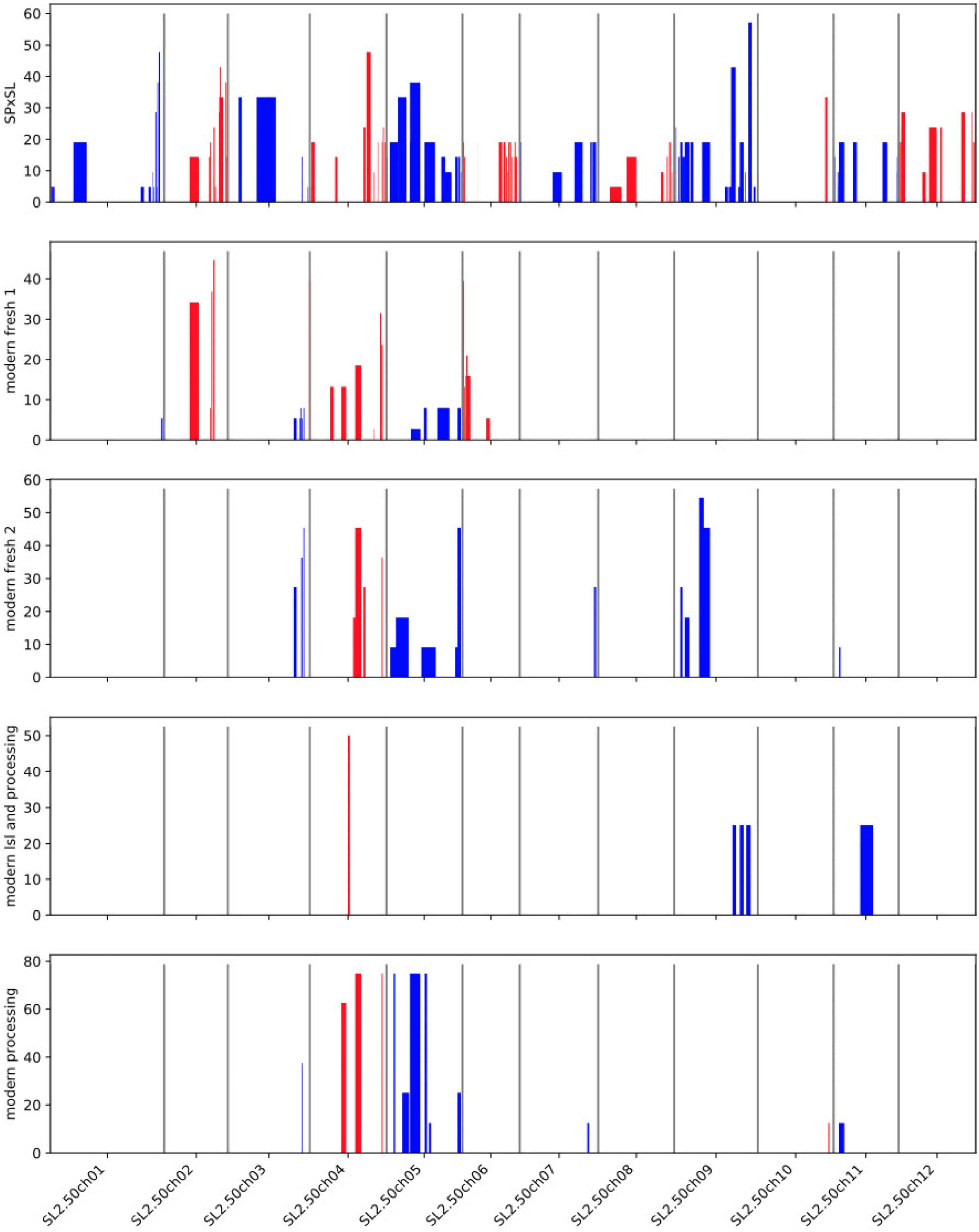

SPxSL
modern fresh 1
modern fresh 2
modern lsl and processing
modern processing
SL2.50ch01
SL2.50ch02
SL2.50ch03
SL2.50ch04
SL2.50ch05
SL2.50ch06
SL2.50ch07
SL2.50ch08
SL2.50ch09
SL2.50ch10
SL2.50ch11
SL2.50ch12

Fig. S3. Introgressed regions along the genome detected in the groups classified as modern: modern flesh 1, modern flesh 2, modern lsl and processing, modern processing, as well as the admixture SPxSL group. Bars indicate the presence of introgression in the chromosomes depicted bellow. The high of the bars indicate the percentage of accesions within group that carry the introgressions according with the scale on the left. Bar colors are only to facilitate the interpretation of the table, blue for odd and red for even chromosomes.

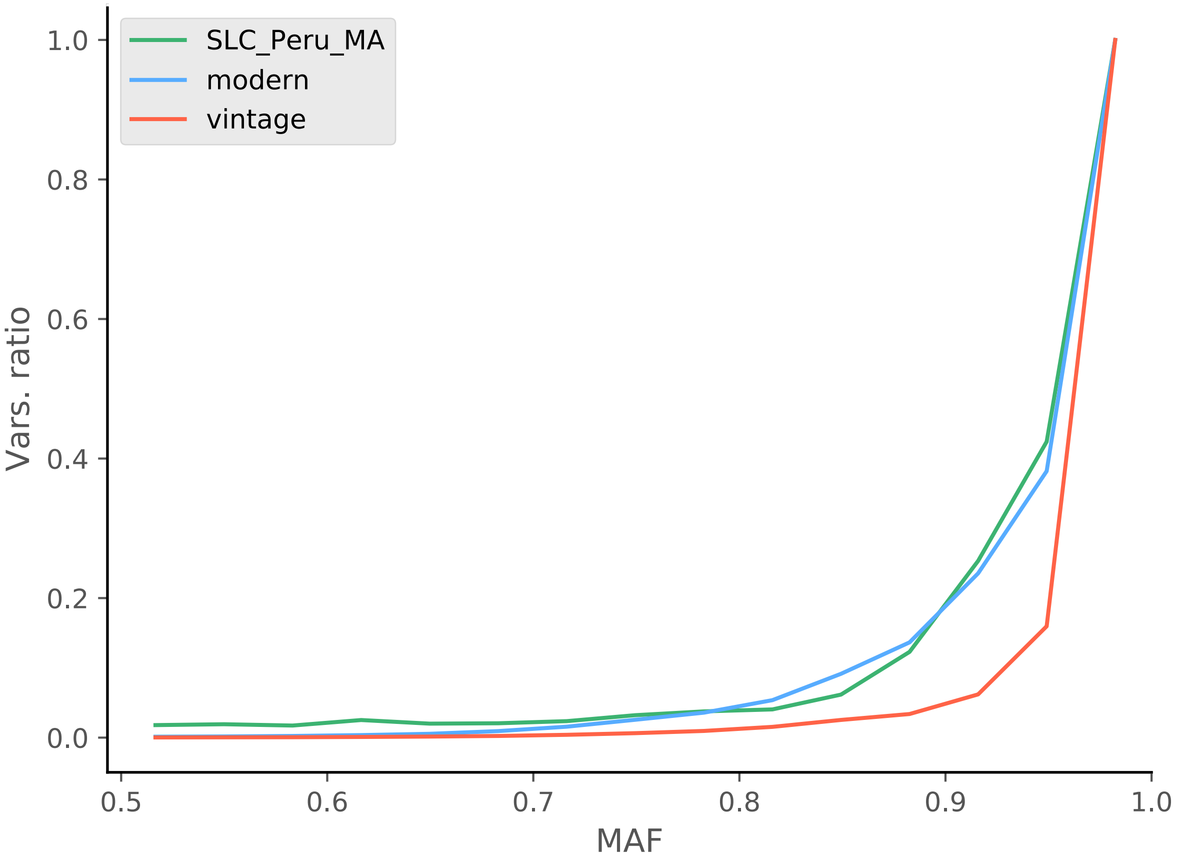

SLC_Peru_MA
modern
vintage
1.0
0.8
0.6
0.4
0.2
0.0
Vars. ratio
0.5
0.6
0.7
0.8
0.9
1.0
MAF

Fig. S4. Major Allele Frequency spectrum in vintage, modern, and SCL_Peru_MA

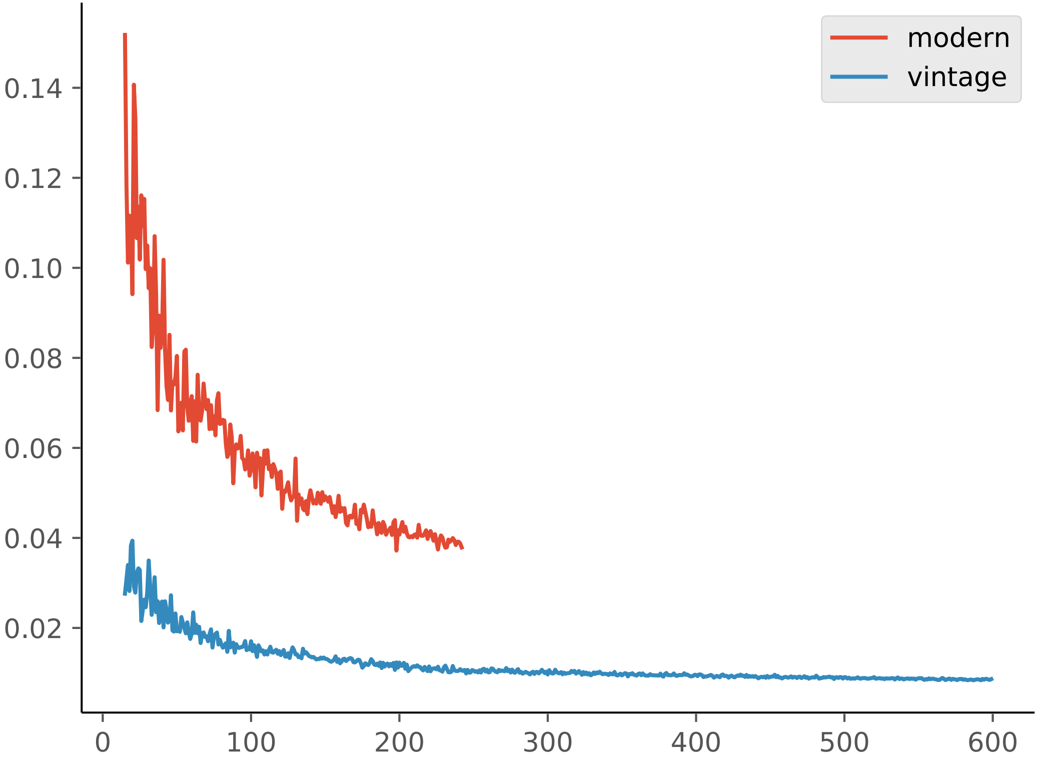
modern
vintage
0.14
0.12
0.10
0.08
0.06
0.04
0.02
0
100
200
300
400
500
600

Fig. S5. Rarefaction analysis of the expected heterozygosity for each genetic group. Axis X shows the number of samples, Axis Y shows the number of variants

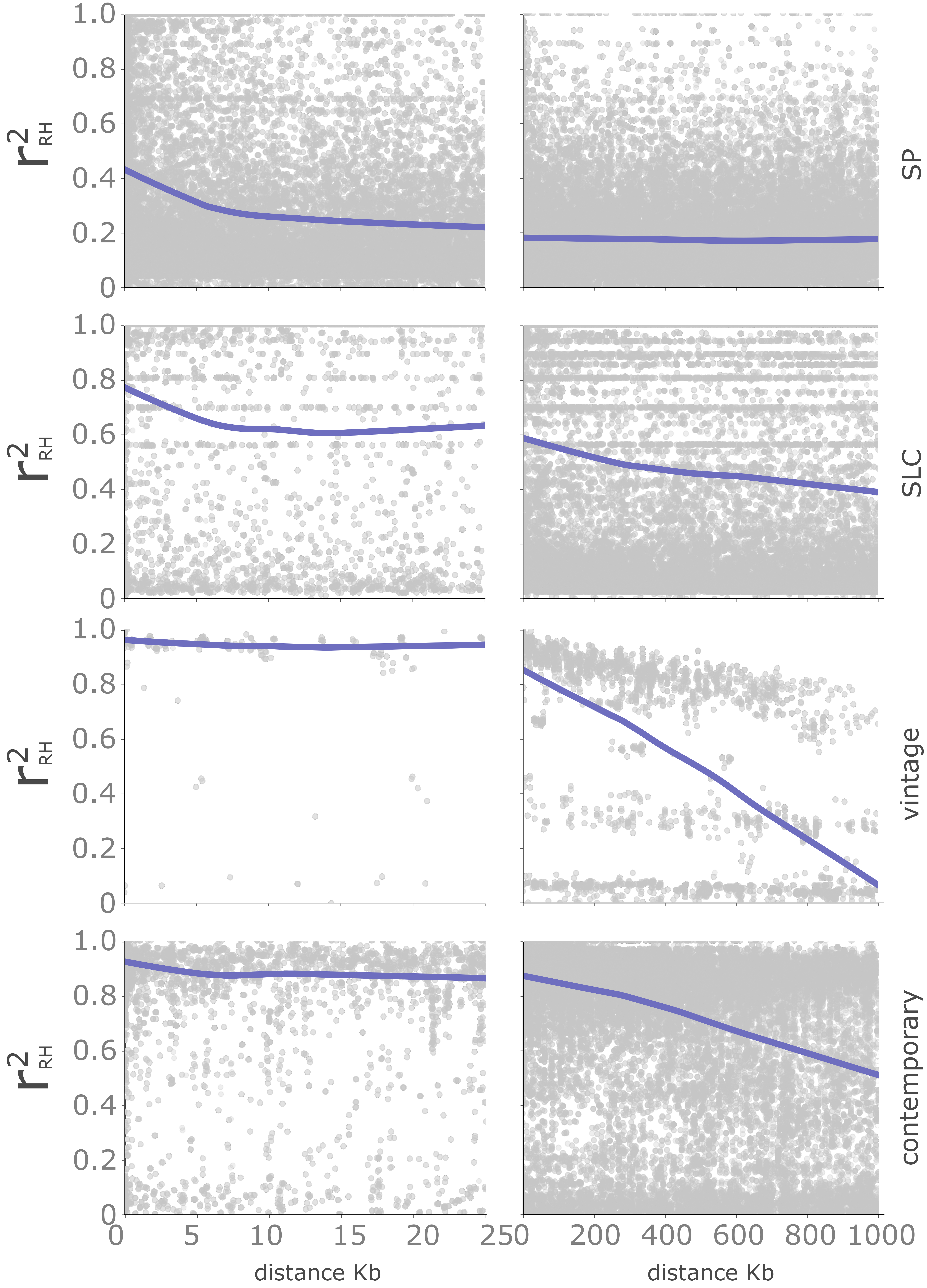

$r^2_{RH}$
SP
SLC
vintage
contemporary
0
0.2
0.4
0.6
0.8
1.0
0
5
10
15
20
250
200
400
600
800
1000
distance Kb
distance Kb

Fig. S6. Genome-wide linkage disequilibrium (LD) decay plot against the physical distance at short (0-25 Kb) and long (0-1,000 Kb) *Solanum pimpinellifolium* (SP), *S. lycopersicum* var. *cerasiforme* (SLC), vintage, and modern accession groups. Linkage disequilibrium (r2) was calculated between euchromatic markers with a major allele frequency lower than 0.98 following the Rogers and Huff (RH) method for loci with unknown phase between pairs of polymorphisms

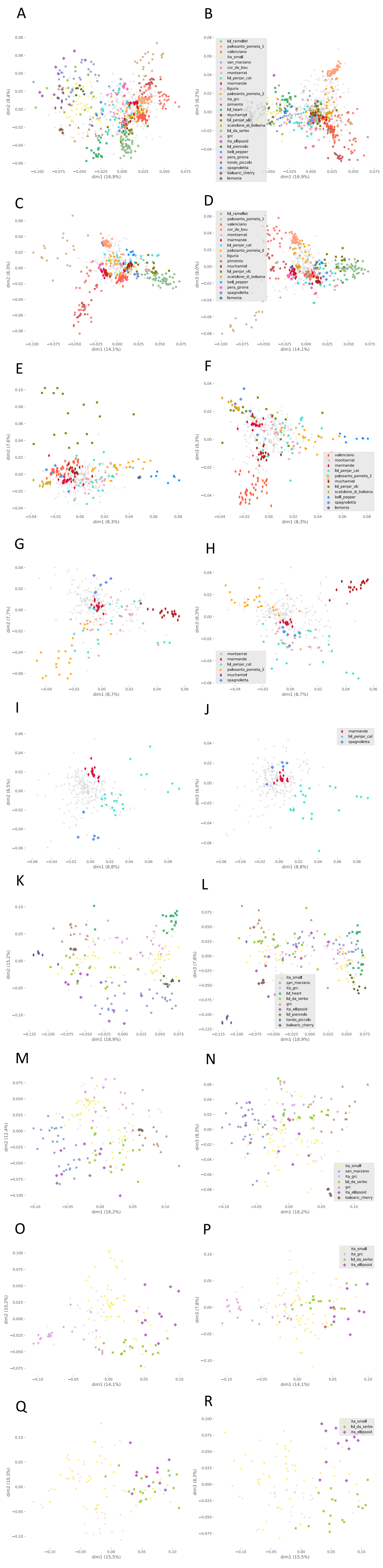

Fig. S7.Hierarchical Principal Coordinate Analysis of European vintage tomato varieties

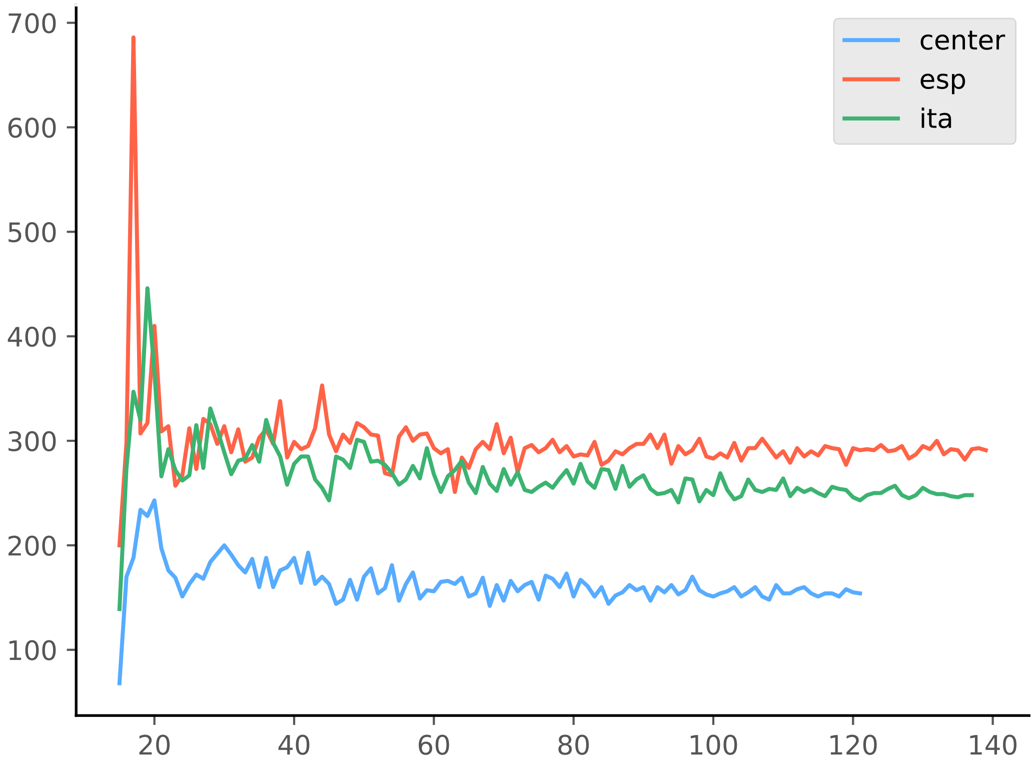

center
esp
ita
700
600
500
400
300
200
100
20
40
60
80
100
120
140

Fig. S8.Rarefaction analysis of the number of polymorphic variants (95% threshold)
